# Spatially Resolved Gene Expression is Not Necessary for Identifying Spatial Domains

**DOI:** 10.1101/2023.10.15.562443

**Authors:** Senlin Lin, Yi Zhao, Zhiyuan Yuan

**Author notes:** Corresponding Author: Yi Zhao Zhiyuan Yuan.

## Abstract

The development of Spatially Resolved Transcriptomics (SRT) technologies has revolutionized the study of tissue organization. We introduce a graph convolutional network with an attention and positive emphasis mechanism, named “BINARY,” relying exclusively on binarized SRT data to delineate spatial domains. BINARY outperforms existing methods across various SRT data types while using significantly less input information. Our study suggests that precise gene expression quantification may not always be essential, inspiring further exploration of the broader applications of spatially resolved binarized gene expression data.

## Main Article

Advances in Spatially Resolved Transcriptomics (SRT) technologies have significantly contributed to the understanding of tissue organization^1,2^. Using SRT data, spatial clustering, akin to single-cell clustering in the field of single-cell RNA sequencing (scRNA-seq), can identify spatial domains in tissues, shifting the focus from a cell-centric understanding of tissue physiology to a higher-level structure-centric perspective^3^. At present, at least 70 spatial clustering methods have emerged, aiming to devise methodologies for modeling gene expression profiles and spatial information to facilitate the identification of spatial domains^4,5^. The current trend in method development involves integrating more information to identify spatial domains^6-8^. For instance, in addition to the two primary modalities of SRT data (i.e., gene expression and spatial coordinates) as commonly done by most methods^9-12^, new methods integrate supplementary images (e.g., immunohistochemistry images^13^, H&E images^14,15^, Chromatin images^16^, etc.), human annotations^17^, paired scRNA-seq data^18^ or other-spatial-omics data^19^. In this context, we explore in an opposite direction, investigating whether it is possible to accurately identify spatial domains with less information. We ask a fundamental question: Is complete spatially resolved gene expression necessary for identifying spatial domains?

Here, we have devised a new spatial clustering method that exclusively relies on binarized SRT data (indicating the presence or absence of gene expression), which we refer to as “BINARY” (Fig. 1A). BINARY is constructed as a graph convolutional network with an attention mechanism^20^. We have also introduced the positive emphasis mechanism by implementing a novel positive-aware binary cross-entropy (PBCE) loss, which penalizes wrong 0s and wrong 1s in an unbalanced manner, rather than merely focusing on the plain reconstruction of binary signals (see Methods). The motivation behind this is the 1s are more credible than 0s in the binary gene expression matrix, due to the nature of spatial transcriptomics data generation process^21,22^. The SRT input data containing the gene expression (GEX) matrix and spatial coordination matrix is encoded into a graph structure, where edges are determined by the spatial relationships between cells, and each node is associated with the binarized gene expression vector (see Methods). Through unsupervised training, BINARY generates embeddings for all nodes within this graph, which are subsequently utilized for clustering to delineate spatial domains and for color space embedding to visualize tissue landscape^23^ (see Methods). BINARY’s workflow is illustrated in Figure 1A. With purpose-built models designed to handle binary data effectively, BINARY eliminates the need for SRT data preprocessing steps (currently still controversy^21^) like normalization and logarithmic transformations, as previous methods did^12,24^. It offers other advantages, such as lower demands on the mRNA capture rate of experimental techniques and the potential to extend model scalability beyond current methods.

**Fig. 1.**
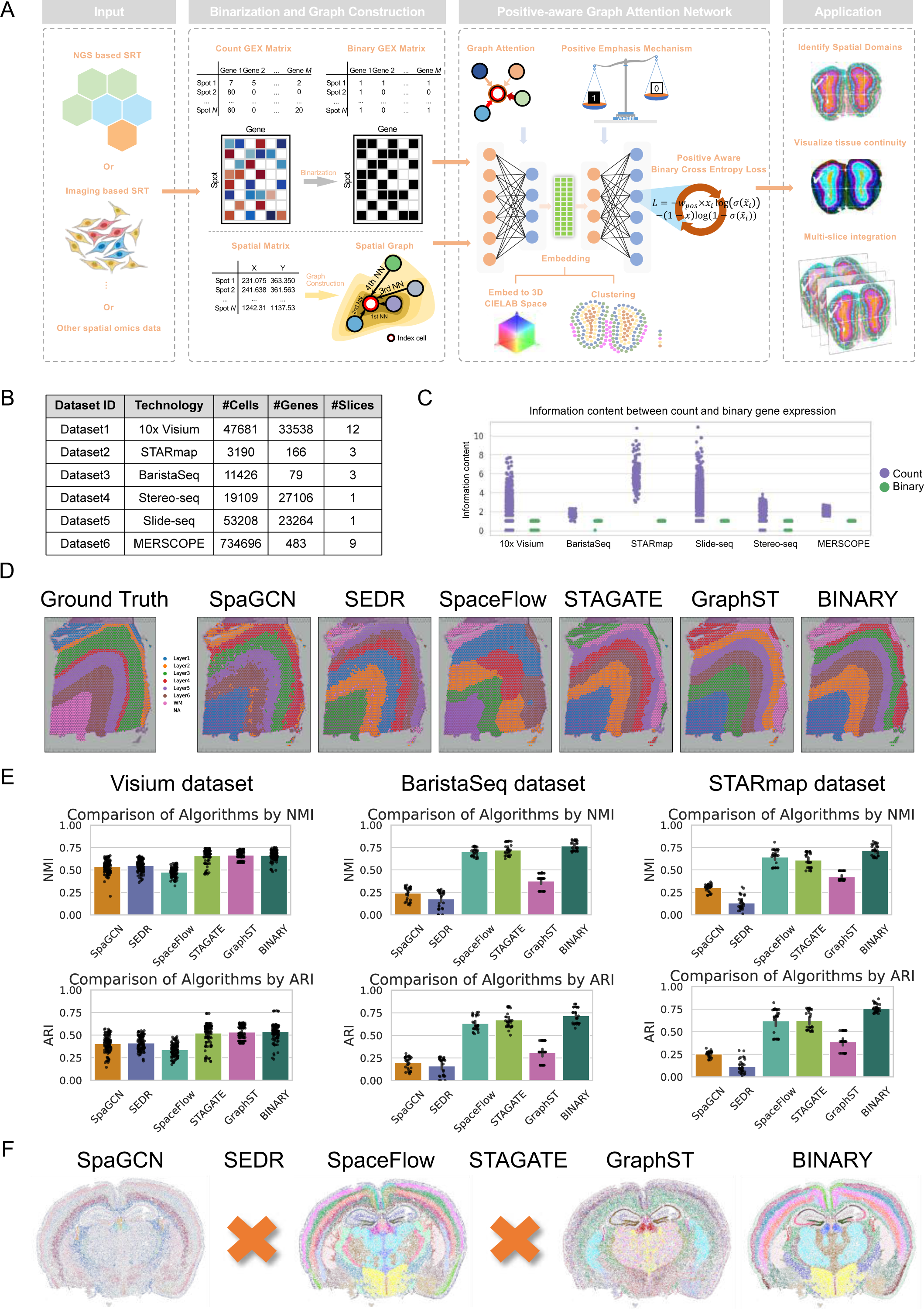
Workflow of BANARY and benchmarking evaluation. **a,** Schematic representation of the BINARY process for the identification of spatial domains using binarized spatially resolved transcriptomics (SRT) data. (i) The procedure initiates with input data generated from various spatial technologies such as next-generation sequencing (NGS) based SRT, imaging-based SRT, or other spatial omics data. (ii) The spatial data contains a gene expression (GEX) matrix and a spatial matrix. In the GEX matrix, rows represent cells (or spots), and columns represent genes. In the spatial matrix, rows represent cells/spots, and columns represent x/y coordinates. The subsequent phase involves a binarization procedure that transforms the raw count GEX matrix into a binary format, and constructs a spatial graph using the spatial matrix. (iii) Following this, BINARY employs the Positive-Aware Graph Attention Network for training the embeddings of cells/spots, which is a lower-dimensional representation of the binary expression data that preserves the essential topological and contextual information. The embedding is used for color space embedding and clustering. (iv)The applications of BINARY include the identification of spatial domains, visualization of tissue continuity, and integration across multiple slices. **b,** Spatial transcriptomics datasets for benchmarking evaluation including their id, technologies and the number of cells (or spots), genes (or features) and slices. **c,** Evaluation of information content across various spatial transcriptomics platforms, binary gene expression data contains substantially less information compared to the unaltered count data. Each point is a gene, the y-axis quantifies how many binary bits are needed to represent the information for each gene. **d,** The clustering results of spatial domains by SpaGCN, SEDR, SpaceFlow, STAGATE, GraphST and BINARY on slice 151673 of the DLPFC dataset. Complete clustering results of the other DLPFC slices are shown in Supplementary Fig. 1. **e,** Quantitative Analysis on Visium, BaristaSeq, and STARmap Datasets. Left: Evaluation of six algorithms using NMI (top) and ARI (bottom) for the Visium dataset (DLPFC, 12 slices). For a comparative visual representation of clustering outcomes and detailed quantitative results across the 12 slices by these six methods, refer to Supplementary Fig 1 and Supplementary Fig 2. Middle: Performance assessment of six algorithms via NMI (top) and ARI (bottom) metrics for the BaristaSeg dataset (3 slices). For a detailed visual representation of clustering results and quantitative insights for each slice by these methods, consult Supplementary Fig 3. Right: Comparative analysis of six algorithms employing NMI (top) and ARI (bottom) metrics for the STARmap dataset (3 slices). For a visual overview of clustering outcomes and quantitative evaluation for each slice by these algorithms, see Supplementary Fig 4. Each point in three barplots signifies an individual trial. Every trial consists of ten runs for each slice across the datasets. **f**, Comparative clustering performance of various state-of-the-art methods on large-scale MERSCOPE data (R3S2 as a representative slice). The orange cross symbols indicate methods, such as SEDR and STAGATE, that encountered ‘out of memory’ errors. The clustering outcomes of the other four methods across nine slices can be referenced in Supplementary Fig 7. For insights on BINARY’s visualization of tissue continuity across the nine MERSCOPE slices, please refer to Supplementary Fig 14.”

We conducted a comprehensive benchmarking study comparing BINARY with recent state-of-the-art spatial clustering methods, which included SpaGCN^15^, SEDR^25^, STAGATE^24^, SpaceFlow^12^, and GraphST^26^, across 6 Spatially Resolved Transcriptomics (SRT) technologies (Fig. 1B). These spatial technologies span a spectrum from next-generation sequencing (NGS)-based to imaging-based technologies^5^. Notably, unlike BINARY, all the other methods make use of complete gene expression count data (SpaGCN incorporated additional histological images), which contains significantly more information than the binary gene expression data (Figure 1C). On the widely used NGS-based 10x Visium human dorsolateral prefrontal cortex (DLPFC) dataset^27^, BINARY delineated the laminar structures with a similar quality compared to recent methods (STAGATE and GraphST), and outperformed the other three methods (SpaGCN, SEDR, and SpaceFlow). This is evident both visually (Fig. 1D, Supplementary Fig. 1) and quantitatively in terms of normalized mutual information (NMI) and adjusted rand index (ARI) (Fig. 1E, 1st column, Supplementary Fig. 2). Imaging-based SRT technologies generate data with very different data characteristics compared to NGS-based data. The most notable difference is that imaging-based SRT data generally exhibit lower gene throughput and a higher mRNA capture rate than NGS-based data^28^. Considering this, we also compared BINARY with other methods on two imaging-based datasets, including BaristaSeq^29^ and STARmap^30^ (Fig. 1B). The results demonstrate that BINARY outperforms other competing methods (Fig. 1E, 2nd and 3rd columns, and Supplementary Fig. 3-4). The above comparison indicates that BINARY utilizes SRT information more efficiently than other methods, irrespective of the trade-offs between mRNA capture rate and gene throughput in the data. Extensive analyses confirm BINARY’s robustness to hyperparameters (Supplementary Fig. 8-10).

Furthermore, BINARY was tested to accurately identify the expected tissue structures in datasets considered more complex, such as the Slice-seq Hippocampus data^31^ (Fig. 1B, Supplementary Fig. 5) and the Stereo-seq Olfactory bulb data^32^ (Fig. 1B, Supplementary Fig. 6). The spatial visualization of the color-coded BINARY embedding can reflect the tissue continuity, even the binary discretization was adopted to the raw SRT data (Supplementary Fig. 13). In an even more challenging dataset containing 9 whole brain sections with more than 730,000 cells and more complex tissue structures, BINARY and SpaceFlow excelled in refining the complex brain structure (BINARY obtained even clearer brain structures than SpaceFlow) compared to other methods (Fig. 1F, Supplementary Fig. 7). It’s worth noting that two of the other methods (SEDR and STAGATE) couldn’t produce results due to excessive memory usage (NVIDIA A100 Tensor Core GPU, 80GB, see Methods). The color-coded visualization is also shown in Supplementary Figure 14. This represents the first demonstration that using only the binary expression of 483 spatially resolved genes, the complex brain structure can be automatically defined at single-cell resolution^33,34^.

With the advances in high-throughput spatial technologies, spatially resolved measurements across various samples/slices at scale are increasingly accessible^35-38^. Multi-slice analysis holds paramount importance in the context of multi-slice and multi-sample experimental designs in modern spatial biology studies as it facilitates the generation of aligned and comparable spatial domain labels across different slices^39^. We have devised a multi-slice spatial graph construction strategy, which has been implemented in BINARY, allowing for joint BINARY training on multiple slices (Fig. 2A, see Methods). We attempted to test the integration analysis of two MERSCOPE brain data using all the competing methods and found that only STAGATE, GraphST, and BINARY provided integration function in their codebase. Among these, only BINARY could run the analysis without exceeding the memory limit (Fig. 2B). By referring to the known brain anatomical structure (Fig. 2C, obtained from Ref^36^), BINARY can not only identify known brain structures but also ensure that label assignments are automatically aligned between the integrated slices, indicating the benefit of multi-slice analysis (Fig. 2D-E).

**Fig. 2.**
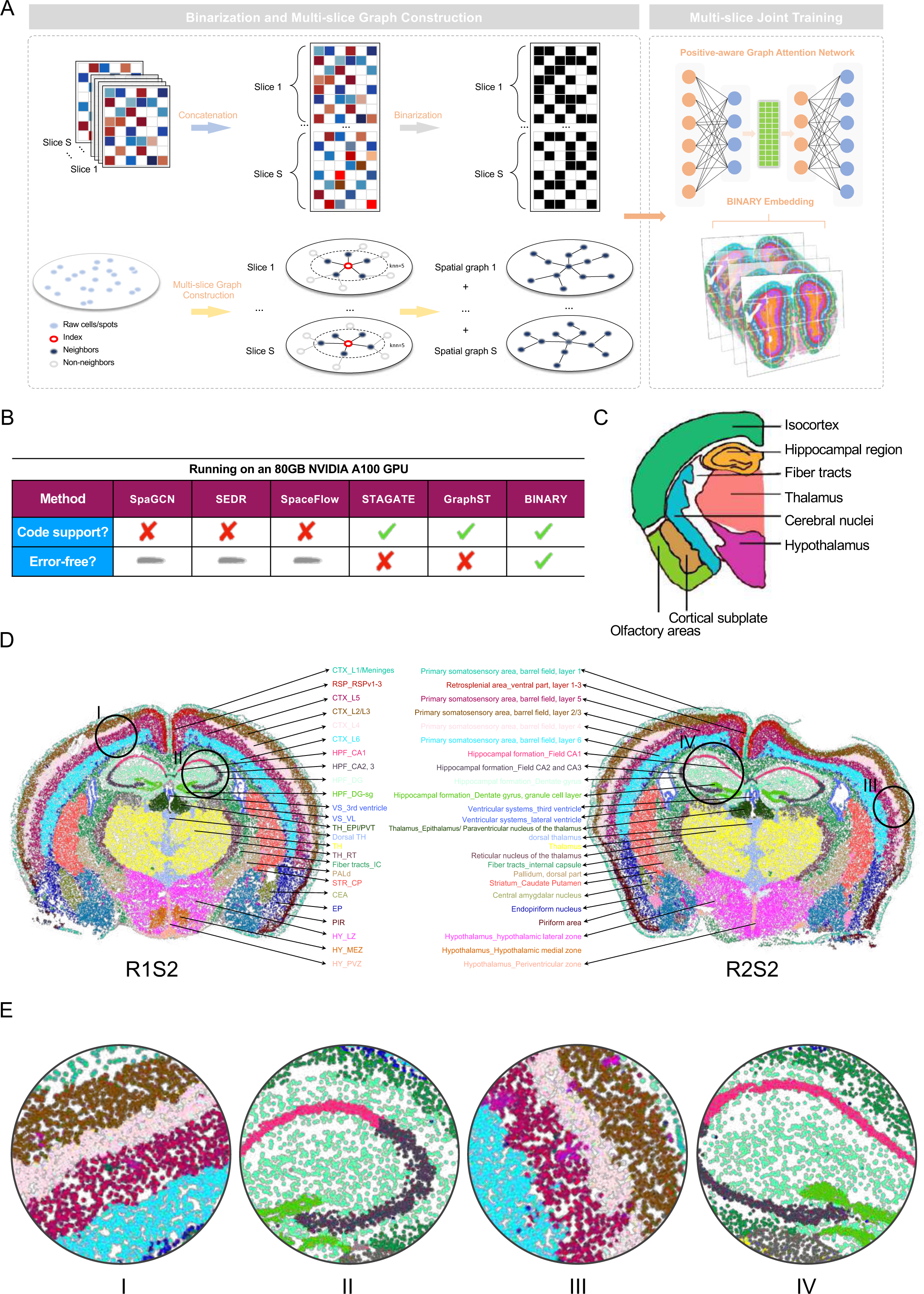
BANARY enable multi-slice analysis on the large-scale multi-slice dataset. **a,** The pipeline for multi-slice joint training: BINARY concatenates data from various slices and proficiently segregates distinct slice-specific raw cells/spots, minimizing potential confusion arising from overlapping spatial information, to facilitate multi-graph construction. Subsequently, they act as inputs for the positive-aware graph attention network. The BINARY’s embeddings are leveraged to multi-slice integration through multi-slice joint training. **b,** BANARY is the sole method capable of supporting multi-slice analysis on a large-scale multi-slice dataset, running seamlessly on an 80GB NVIDIA A100 GPU. Other methods either lack the requisite code design or encounter memory errors. **c,** Mouse Brain Anatomy Reference. **d,** Multi-slice joint training yields consistent clustering identifications across MERSCOPE slices. **e,** Detailed local zoom-in visualizations of specific regions (I, II, III, IV) from the slices presented in (d).

This study goes beyond the development of a novel computational method; it also motivates a rethinking in the field: for the identification of spatial domains, precise gene expression quantification through experimental technologies in spatial transcriptomics may not be necessary, while a binary value indicating the presence or absence of gene expression suffices. These findings have the potential to reshape the landscape of spatial clustering and other computational tasks in the study of spatial biology within the field of spatial transcriptomics and other spatial omics (as demonstrated by a BINARY application on two spatial proteomics data in Supplementary Fig. 12).

## Methods

### BINARY

### Binarization

We propose, for the first time, the potential of substituting binary expression matrices in place of count-based expression matrices in spatial transcriptomics. This entails directly utilizing the binary representation of spatial transcriptomics data rather than its count version. Before training, we process the original SRT count data, denoted as X_*count*_, through binarization processing. This results in the binary expression matrix for spatial transcriptomics, denoted as X_*binary*_:

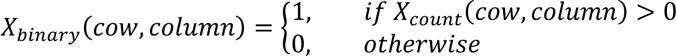

where the row and column are the indices for cell/spot and gene. Note that the original count matrix can be either subset of whole transcriptome (demonstrated using imaging-based SRT data) or suffering from low mRNA capturing rate (demonstrated using NGS-based SRT data).

### Spatial graph construction

The primary advantage of spatial transcriptomics lies in its ability to capture both spatial information and gene expression of cells/spots. To emulate the topological associations among these spots or cells, spatial graph construction emerges as a standard representation. Constructing spatial graphs can enable models to leverage the underlying structure, allowing the extraction and learning of feature expressions from neighboring spots/cells. In the process of constructing spatial graphs, the Construct_Spatial_Graph() function presents two methodologies: the K-Nearest Neighbors (KNN) and the Radius-Based approaches. Using the KNN method, each cell/spot, is linked to its ‘k’ closest neighbors based on spatial coordinates. Here, ‘cutoff’ aligns with ‘k’. Conversely, the Radius-Based approach connects each cell/spot to all other entities within a predetermined radius, denoted by ‘cutoff’. The KNN method is our default approach for spatial graph construction, primarily due to the inherent challenges associated with controlling the number of neighbors for each cell or spot when employing the radius-based approach. When modifying the radius ‘cutoff’, there can be substantial variance in neighbor counts for cells or spots across different datasets. To elaborate, irrespective of whether one uses the KNN or the radius-based method, for the 10 x Visium dataset, it’s advisable to keep the number of neighbors for each spot between 5 to 8. A comprehensive sensitivity analysis concerning the selection of neighbor parameters across different datasets can be found in Supplementary Fig 8, Supplementary Fig 9, and Supplementary Fig 10. Furthermore, to facilitate co-training across multiple slices, we’ve incorporated a feature for multi-slice spatial graph construction. With the Mutil_Construct_Spatial_Graph() function, users can construct graphs for datasets with multiple slices, ensuring each slice maintains its accurate spatial relationship. This is crucial for subsequent multi-slice joint training, as depicted in Fig 2A.

### Positive-aware graph attention network

To utilize the binarized expression matrix and the spatial graph, we meticulously craft a positive-aware graph attention network for identifying tissue domains. Building upon the graph auto-encoder, the positive-aware graph attention network integrates a graph attention mechanism that takes into account self-features, along with a novel positive emphasis mechanism. Ultimately, the entire model is optimized with a specialized positive-aware binary cross-entropy loss function, explicitly designed for binarized expression data.

### Graph attention

The graph attention allows nodes to focus on their neighbors with varying weights, based on both their own features and those of their neighbors. These attention weights are computed through a learnable adaptive mechanism. Generally speaking, the attention weights are derived from the features of the node and its neighboring nodes. For every pair of cells or spots, its graph attention score *e_ij_* can be derived as:

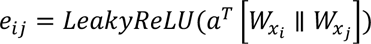

where ∥ denotes the concatenation operation, *a* is the weight for graph attention, which is a trainable vector. *W* is a weight matrix that serves the purpose of translating the input features of cells or spots into a new representational space. For every cell or spot, all attention coefficients are normalized using the softmax function to ensure their summation equals 1:

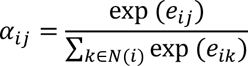

where *N*(*i*) denotes the neighbors of cell or spot *i*. In the first layer of the graph auto-encoder considering the graph attention, the node’s own features are also taken into account. This is achieved by adding a self-loop to the graph, ensuring that its features are considered when aggregating features from neighboring cells or spots.

### Positive emphasis mechanism

Given the intrinsic sparsity of the original count matrix and the subsequent binarization, positive entries (represented as “1”) in the binarized expression matrix encapsulate particularly vital information. Such entries demand augmented attention during the model’s training phase, ensuring that these pivotal data points are effectively captured and represented. To address this, we introduced the positive emphasis mechanism (PEM). PEM quantitatively translates the importance of positive entries into a weight parameter, denoted as *w*_*pos*_. This weight is seamlessly integrated with the loss function.

### Graph auto-encoder

A graph auto-encoder (GAE) architecture is employed to learn latent representations from binarized expression matrices. Specifically, the encoder captures the low-dimensional embeddings (hidden layer) of the binarized data suitable for downstream tasks, while the decoder aims to reconstruct the initial binarized signals. Our adaptation of GAE introduces two pivotal enhancements: (i) Graph attention: This refinement allows the model to weigh the importance of nodes based on their intrinsic features and their relationship with neighboring nodes. (ii) Positive emphasis mechanism: Utilizing the intrinsic sparsity in the binarized expression data, this mechanism amplifies the significance of positive signals in the learning process. For the optimization of model parameters, we employ a specialized loss function tailored for binarized expression data, termed the positive-aware binary cross-entropy loss.

The encoder consists of two layers leading to the generation of embeddings from a binarized expression matrix. The network accepts a binarized expression matrix X_*binary*_ ∈ ℝ^*N*×*M*^_0,_ where *N* is the number of spots/cells and *M*_0_ is the dimension genes/features. In the first layer ℎ^(1)^, for each spot/cell *x*_*i*_ ∈ X_*binary*_, *i* ∈ {1,2,3, …, *N*}, influences from its neighboring spots/cells are incorporated, while also factoring in its inherent significance. The first hidden layer can be described as:

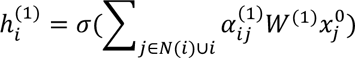

Where *W*^(1)^ ∈ ℝ^*M*_0_×*M*_1_^ denotes the weight matrix for the first layer with *M*_1_ as the hidden dimension. *a_ij_*^(1)^ is the attention coefficient for the first layer and *x*_*j*_^0^represents the initial neighbors of spot or cell *x*_*i*_. Second layer applies a direct linear transformation without the attention mechanism. Given the first layer ℎ^(1)^ and the weight matrix *W*^(2)^ ∈ ℝ^*M*_0_×*M*_1_^, where *M*_2_ is the dimension of the embedding. Second layer is defined as:

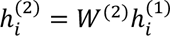

Subsequently, the condensed representation of the binarized matrix, which is appropriate for downstream analyses, is given by:

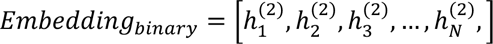

The decoder consists of the third and fourth layers, aimed at reconstructing the binarized matrix representation. The third layer reincorporates the graph attention. Crucially, it adopts a weight matrix transposed from that of the second layer: *W*^(3)^ = (*W*^(2)^)^*T*^. This design facilitates a swifter recreation of the original representation and concurrently aids in mitigating overfitting risks:

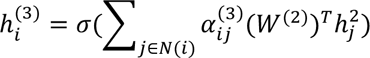

Mirroring the structure of the second layer, this layer merely implements a linear transformation devoid of the graph attention. Using the third layer, ℎ^(3)^, and setting the weight matrix for this layer as *W*^(4)^ = (*W*^(1)^)^*T*^, the output can be stated as:

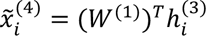

### Positive-aware binary cross-entropy loss function

To ensure the effective reconstruction of positive values, i.e., the value of 1, within sparse binarized expression matrices, a specialized loss function is introduced. Two essential considerations influence the design of this loss function: (i) Binary cross-entropy loss: Given the objective of reconstructing a binarized expression matrix, we employed the binary cross-entropy loss, tailored explicitly for the reconstruction of binary data; (ii) Positive emphasis mechanism: In sparse binarized expression matrices, the occurrence of 1s is much less frequent than 0s. However, these 1s encapsulate a significant amount of information. To guarantee that the model can accurately reconstruct these critical 1s, we introduced a weighting factor, *w*_*pos*_. This ensures that any incorrect prediction regarding the position of a 1 by the model incurs a more substantial penalty, incentivizing the model to prioritize the reconstruction of 1s. The loss function is expressed as:

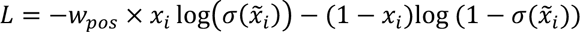

Where *x*_*i*_ represents the true label from the input. *x*X_*i*_denotes the predicted output from the model. σ signifies the activation function, wherein the sigmoid function is employed, mapping the model’s raw output values to the range (0, 1). *w*_*pos*_ is the weight associated with positive emphasis mechanism, with a default value of 10. A comprehensive sensitivity analysis concerning the selection of this parameter across different datasets can be found in Supplementary Fig 8, Supplementary Fig 9, and Supplementary Fig 10.

## Spatial clustering

### Clustering

Upon training the model, the low-dimensional embeddings generated by BINARY are utilized to identify spatial domains. We designed an optimized resolution strategy for Leiden Algorithm. For the Leiden algorithm implemented via the SCANPY package, a method that employs binary search for determining the optimal resolution has been proposed. This algorithm, detailed in Supplementary Fig 11, revolves around the find_optimal_resolution() function. Given a user-defined cluster number, the function leverages binary search to swiftly traverse varying resolution values, culminating in the pinpointing of an optimal resolution that yields the expected cluster number for the given data. The initial resolution range can be user-defined, and the midpoint resolution of the current range is employed for each iteration. Clustering is performed using Leiden. After clustering, the cluster number under the current resolution is computed and juxtaposed with the expected cluster number. If discrepancies arise, the binary search method readjusts the resolution range and adaptive step size, iterating until one of the following conditions is met: identification of a resolution producing the expected cluster number or reaching a predetermined maximum number of iterations. This methodology offers a swifter and more accurate determination of the desired clustering number’s corresponding resolution, especially when compared to SEDR, which demands extended runtimes and utilizes a fixed step size, potentially resulting in failures to identify the correct resolution.

As done by previous spatial clustering methods^15,26^, after the initial clustering outcomes are obtained, an optional step of clustering result optimization can be applied to achieve more coherent and smoother cluster delineations. For any given cell or spot, its label is reassessed and potentially reallocated. This decision is based on the most frequently observed domain among its neighbors within a specified radius ‘r’. By default, r = 30. For multi-slice data, a function tailored for multi-slice refinement, Multi_Refine_label(), is provided. This ensures each slice undergoes individual refinement, mitigating any misalignment issues arising from coordinate overlaps between slices.

### Visualizing tissue Continuity using SOView

In addition to harnessing BINARY’s embeddings for identifying tissue domains via spatial clustering, which presents results with discrete attributes, a more comprehensive representation can be realized to emphasize and visualize the inherent continuity within tissue structures. To achieve this, BINARY’s embeddings can be mapped onto the RGB color space using manifold projection techniques, specifically UMAP (Uniform Manifold Approximation and Projection). Subsequently, this continuous representation can be visualized using the SOView (https://soview-doc.readthedocs.io/en/latest/index.html) tool, providing an overview of spatial organization within tissues.

### Multi-slice joint training

While the prevailing methodologies predominantly focus on individual-slice designs, only a handful, such as STAGATE, are tailored to facilitate multi-slice joint training, specifically for identifying 3D spatial domains. Addressing this limitation, we introduce a more versatile approach termed ‘Multi-slice Joint Training’. This method facilitates the joint training over datasets comprising multiple slices. For an accurate representation of multi-slice datasets, we employed the Mutil_Construct_Spatial_Graph() function. This meticulous method ensures that both intra-slice and inter-slice relationships are preserved and accurately depicted in the spatial graph. Moreover, the concat_adatas() function is invoked to integrate the features of multiple slices seamlessly. This consolidated data forms the substrate upon which multi-slice joint training is executed, as depicted in Fig 2a. The essence of multi-slice joint training lies in its capacity to harness data from individual slices, promoting multi-slice sample learning during model training. This not only promotes mutual information compensation among slices but also ensures consistent annotations across the dataset, as visualized in Fig 2D. Moreover, the proposed multi-slice joint training framework serves as a pivotal reference in the burgeoning era of spatial transcriptomics, accommodating the explosive growth of spatial omics data and providing a unified training reference.

### Comparison with other state-of-the-art methods

In our effort to offer a comprehensive performance overview, we compared our approach with several state-of-the-art methods, specifically: SpaGCN, SEDR, SpaceFlow, STAGATE, and GraphST. The reason for choosing these methods is that they are among the most influential and represent the most recent advanced approaches in this field. While there are many other spatial clustering methods that we cannot test them all, such as BayesSpace^9^, BASS^40^, and SpatialPCA^11^, which also have made significant contributions to the field, we did not include them in our comparison. The primary reason for their exclusion is their lack of GPU acceleration, which could potentially lead to extended running times. To ensure a fair evaluation, instead of uniformly applying default parameters across all datasets, we conducted thorough parameter tuning for each algorithm on each individual dataset to replicate the results as reported in their respective original papers (one can refer to the well-known DLPFC 151673 slice results in Fig. 1D and compare them with the originally reported results in each method’s paper).

## Spatial Data

### Source of Spatial Datasets

All spatial datasets utilized in this study were sourced from the Spatial Omics Database (SODB). The database can be accessed at the following URL: https://gene.ai.tencent.com/SpatialOmics.

### Data Retrieval via Pysodb Python Package

To facilitate data download, we employed the Pysodb Python package. This package provides a seamless interface to the SODB, and its repository can be found on GitHub: https://github.com/TencentAILabHealthcare/pysodb.

### Pysodb Usage Tutorial

For researchers and readers interested in a comprehensive tutorial on how to use the Pysodb package for data retrieval, a detailed guide is available at: https://protocols-pysodb.readthedocs.io/en/latest/.

## Data Processing

BINARY incorporates purpose-built models designed to handle binary data effectively, eliminating the need for preprocessing steps like normalization and logarithmic transformations. Only binarized expression matrix is required as input. When confronted with datasets characterized by a high gene or feature dimensionality, such as the DLPFC dataset that encompasses 33,538 genes, optimization measures can be taken. For improved training efficiency, top 2,000 highly variable genes (HVG) can be selected as inputs for BINARY. Note that the HVG step is performed after the binarization step, ensuring that both data processing and model training step cannot see the original complete gene expression data.

## Computational resource

All experiments are performed on a server running Ubuntu with an Intel(R) Xeon(R) Platinum 8375C CPU *W* 2.90GHz, 120 GB of RAM, and one NVIDIA A100(80G) GPU.

## Reporting Summary

Further information on research design is available in the Nature Research Reporting Summary linked to this article.

## Data availability

Dataset1: http://research.libd.org/spatialLIBD

Dataset2: http://starmapresources.com/data

Dataset3: https://spacetx.github.io/

Dataset4: https://github.com/JinmiaoChenLab/SEDR_analyses/tree/master/data

Dataset5: https://singlecell.broadinstitute.org/single_cell/study/SCP815/

Dataset6: https://vizgen.com/

All datasets are also available at SODB^23^ and can be loaded by pysodb (see the online tutorial in “**Code availability**”).

## Code availability

The Python implementation of BINARY is available at https://github.com/senlin-lin/BINARY/. We also uploaded the code with installation tutorials in the manuscript tracking system. A tutorial on BINARY package is available at https://binary-tutorials.readthedocs.io/en/latest/index.html.

## Acknowledgments

Z.Y. acknowledges the support by Shanghai Municipal Science and Technology Major Project (No.2018SHZDZX01), ZJ Lab, Shanghai Center for Brain Science and Brain-Inspired Technology, and 111 Project (No.B18015). We thank Yining Hao and Weishi Liu for the brain region annotation.

## Author contributions

Z.Y. and S.L. conceived and designed the study, developed the computational methods, performed the analysis, and wrote the manuscript. Y.Z. supervised the work, reviewed the manuscript, and provided important guidance for the work.

## Inclusion & Ethics

Not relevant.

---

**Supplementary Fig. 1.**
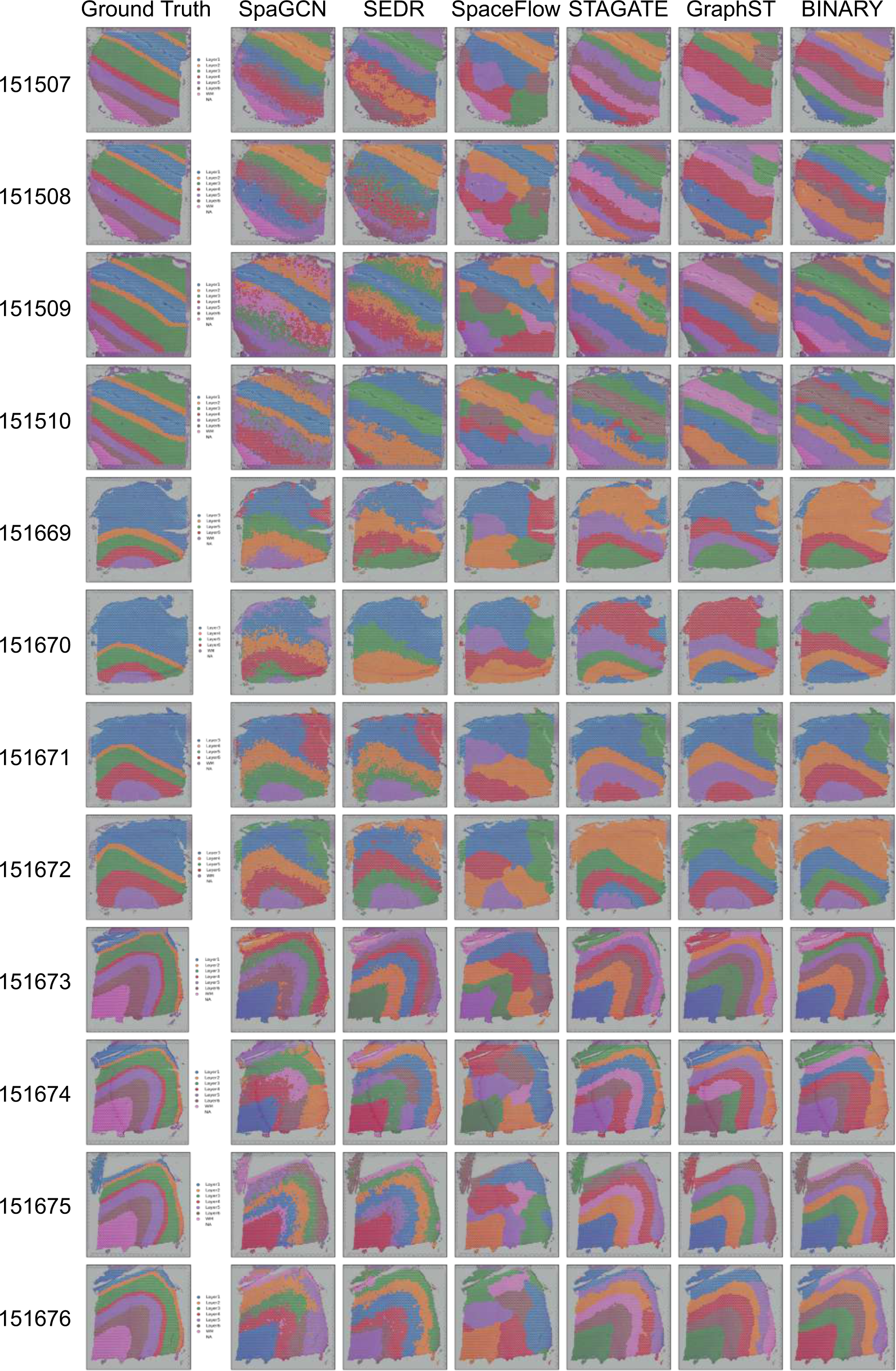
Comparing state-of-the-art spatial clustering methods on DLPFC Visium dataset. Manual annotations and comparison of spatial domains identified by SpaGCN, SEDR, SpaceFlow, STAGATE, GraphST and BINARY on the Dorsolateral Prefrontal Cortex (DLPFC) Visium dataset comprising 12 Slices.

**Supplementary Fig. 2.**
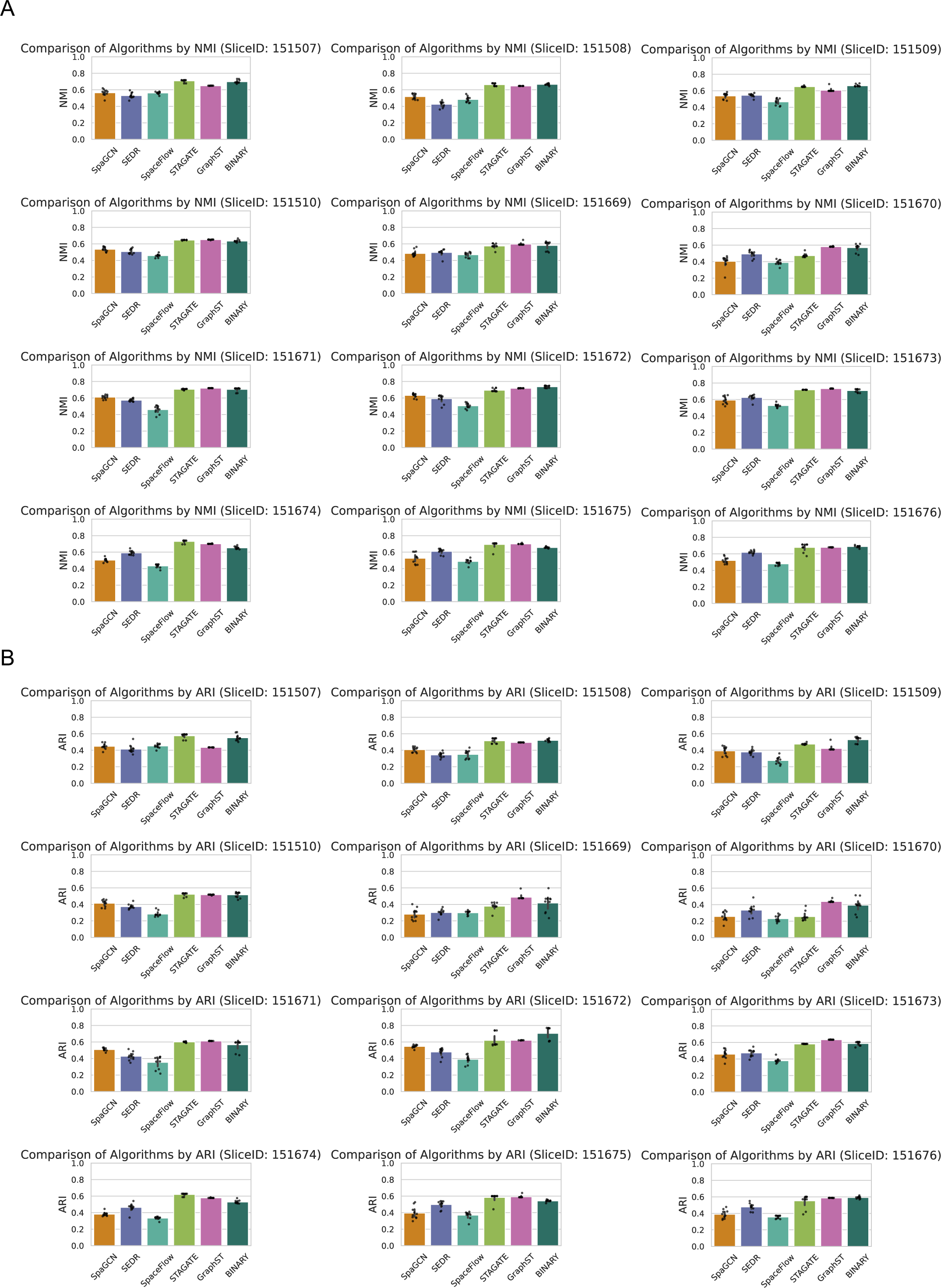
Comprehensive quantitative evaluation of DLPFC Visium dataset using leading spatial clustering approaches. **a,** Comparative analysis using Normalized Mutual Information (NMI) across 12 individual slices. The methods compared include SpaGCN, SEDR, SpaceFlow, STAGATE, GraphST, and BINARY. **b,** Detailed assessment based on Adjusted Rand Index (ARI) across the same 12 slices, using the aforementioned algorithms. Each data point in all barplots represents a distinct experimental trial. For every trial, a series of ten repeated runs was conducted for each slice within the datasets to ensure robustness and reliability of the results.

**Supplementary Fig. 3.**
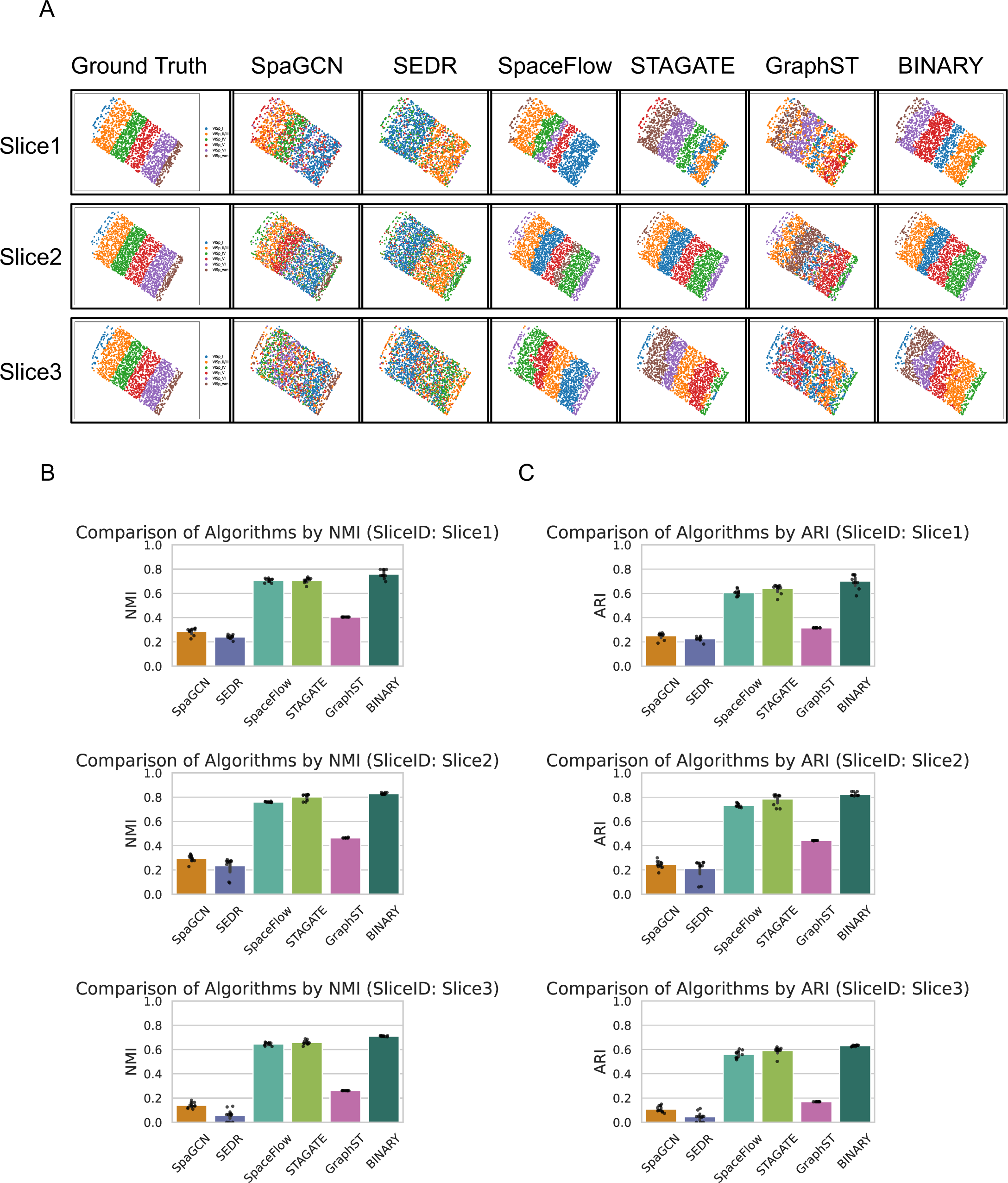
Benchmark evaluation of six methods on the BaristaSeq dataset. **a,** Detailed visual representation showcasing the clustering outcomes for the 3 tissue slices from the BaristaSeq dataset, analyzed using spatial clustering algorithms: SpaGCN, SEDR, SpaceFlow, STAGATE, GraphST, and BINARY. **b,** Quantitative evaluation highlighting the Normalized Mutual Information (NMI) values obtained from the 3 slices across SpaGCN, SEDR, SpaceFlow, STAGATE, GraphST, and BINARY. **c,** Quantitative metrics showcasing the Adjusted Rand Index (ARI) values for the 3 slices, analyzed using the aforementioned spatial clustering methods. Each datapoint depicted in three barplots corresponds to a distinct experimental trial. Notably, every individual trial encompasses ten iterative runs executed on each slice within the complete dataset.

**Supplementary Fig. 4.**
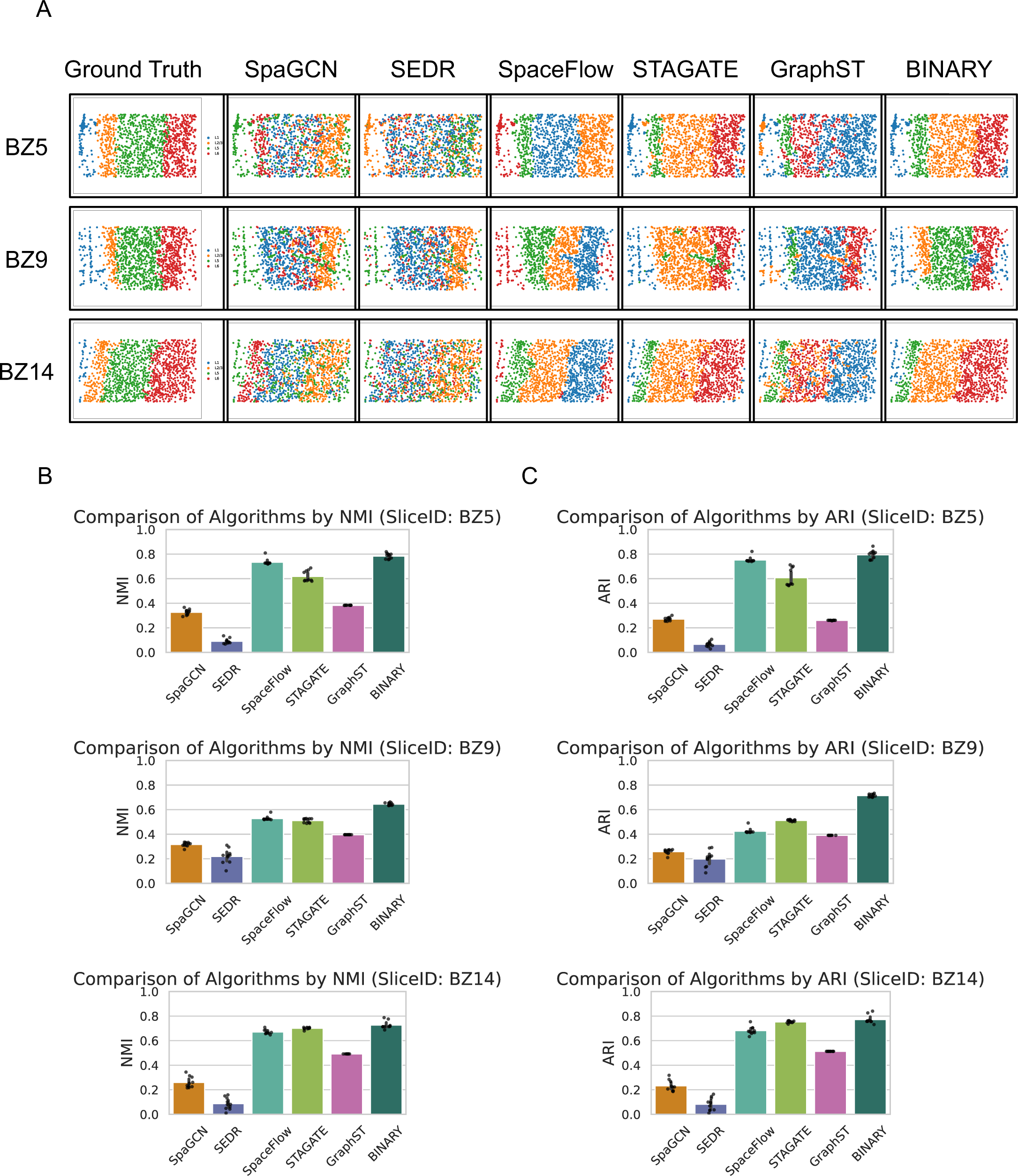
Benchmarking assessment of six methods on the STARmap dataset. **a,** Comprehensive visual depiction of clustering results on the 3 slices STARmap dataset, as analyzed by SpaGCN, SEDR, SpaceFlow, STAGATE, GraphST, and BINARY. **b,** Quantitative evaluation of Normalized Mutual Information (NMI) metrics across the three slices, employing methods such as SpaGCN, SEDR, SpaceFlow, STAGATE, GraphST, and BINARY. **c,** Quantitative comparison of Adjusted Rand Index (ARI) across the three slices. Each data point in the barplots represents an individual trial. Each trial encompasses ten iterative runs for every slice within the dataset.

**Supplementary Fig. 5.**
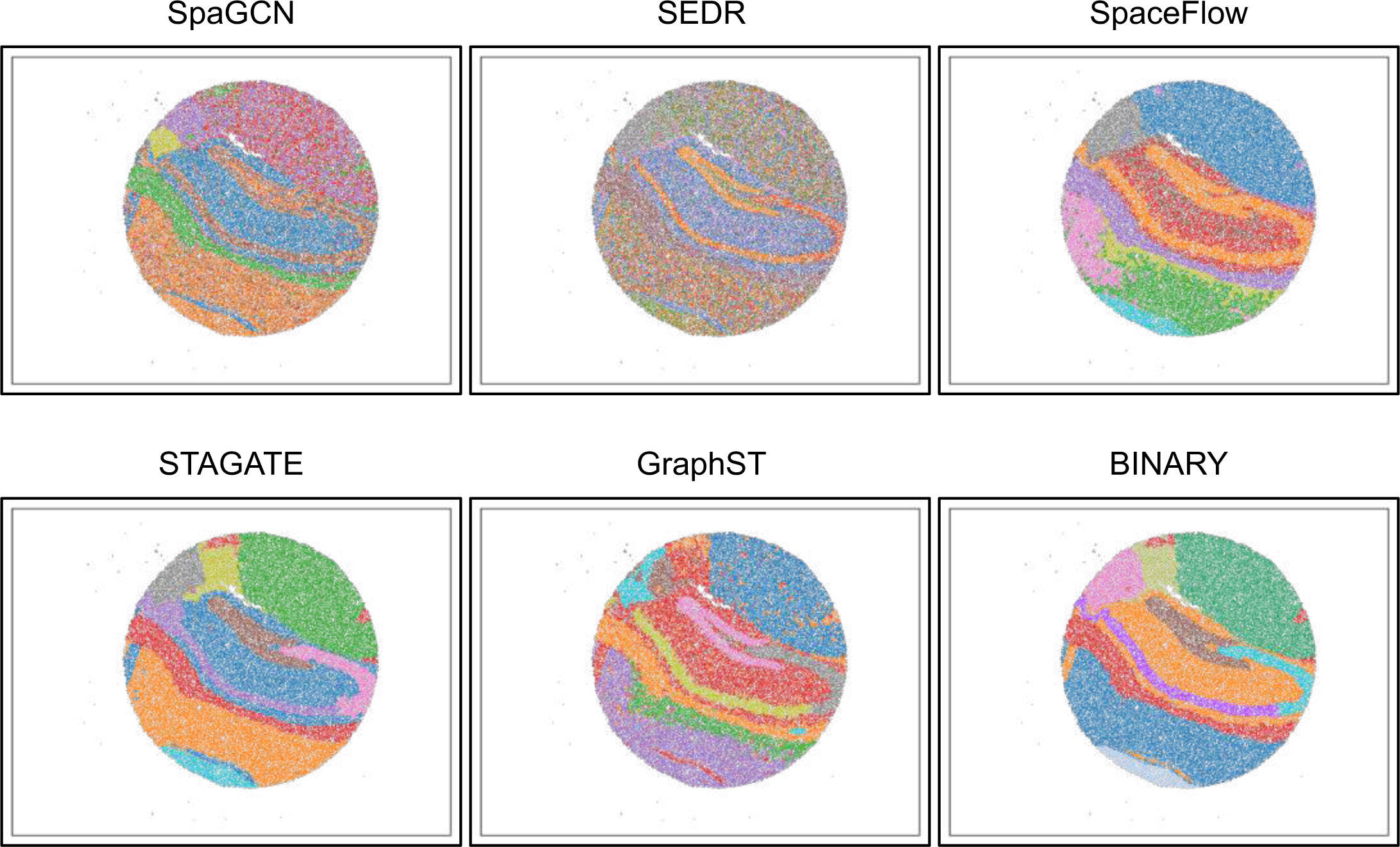
Comparative analysis of spatial domains identified by SpaGCN, SEDR, SpaceFlow, STAGATE, GraphST, and BINARY using the Slide-seq Dataset.

**Supplementary Fig. 6.**
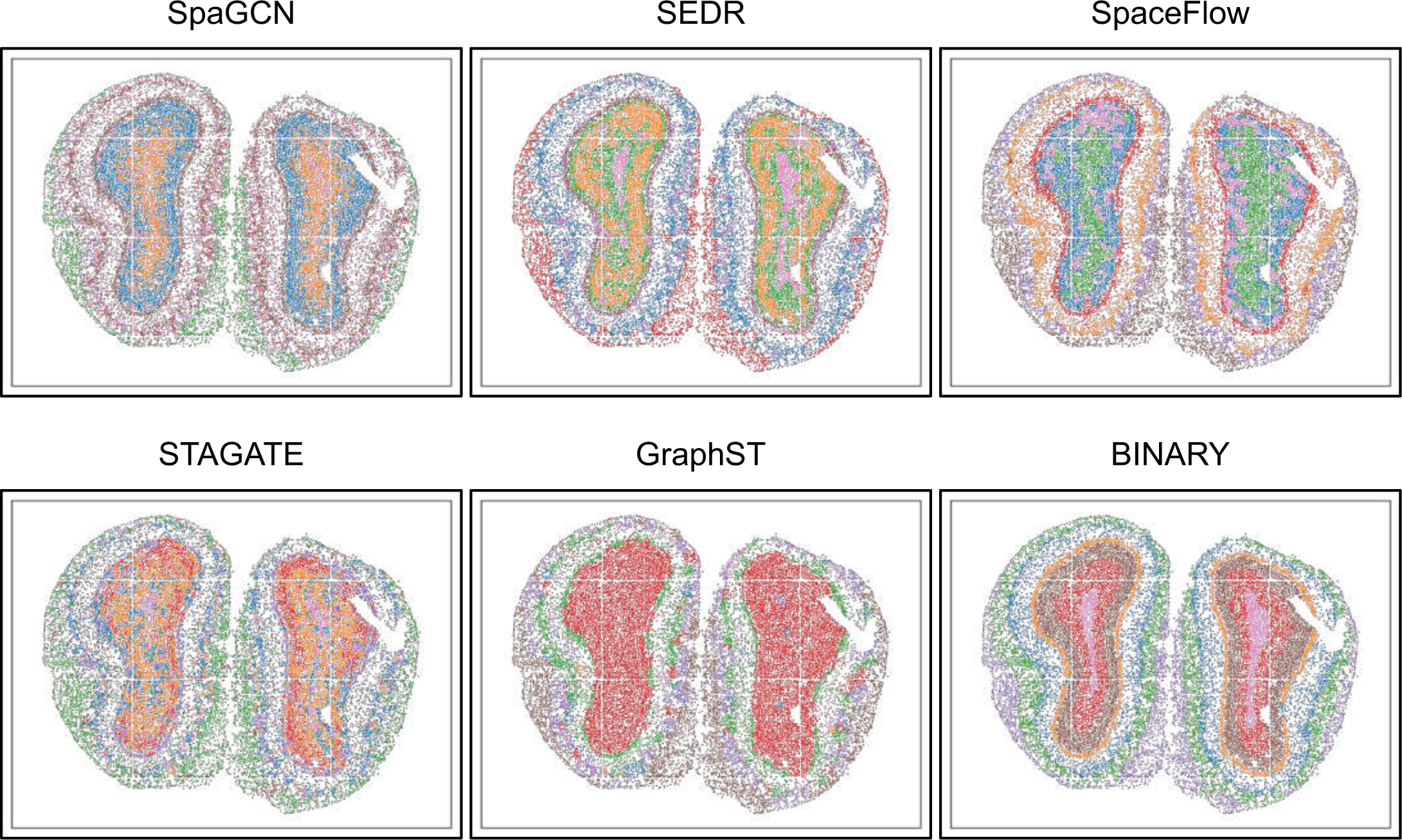
Evaluation of spatial clustering performance on the Stereo-seq dataset by six state-of-the-art algorithms: SpaGCN, SEDR, SpaceFlow, STAGATE, GraphST, and BINARY.

**Supplementary Fig. 7.**
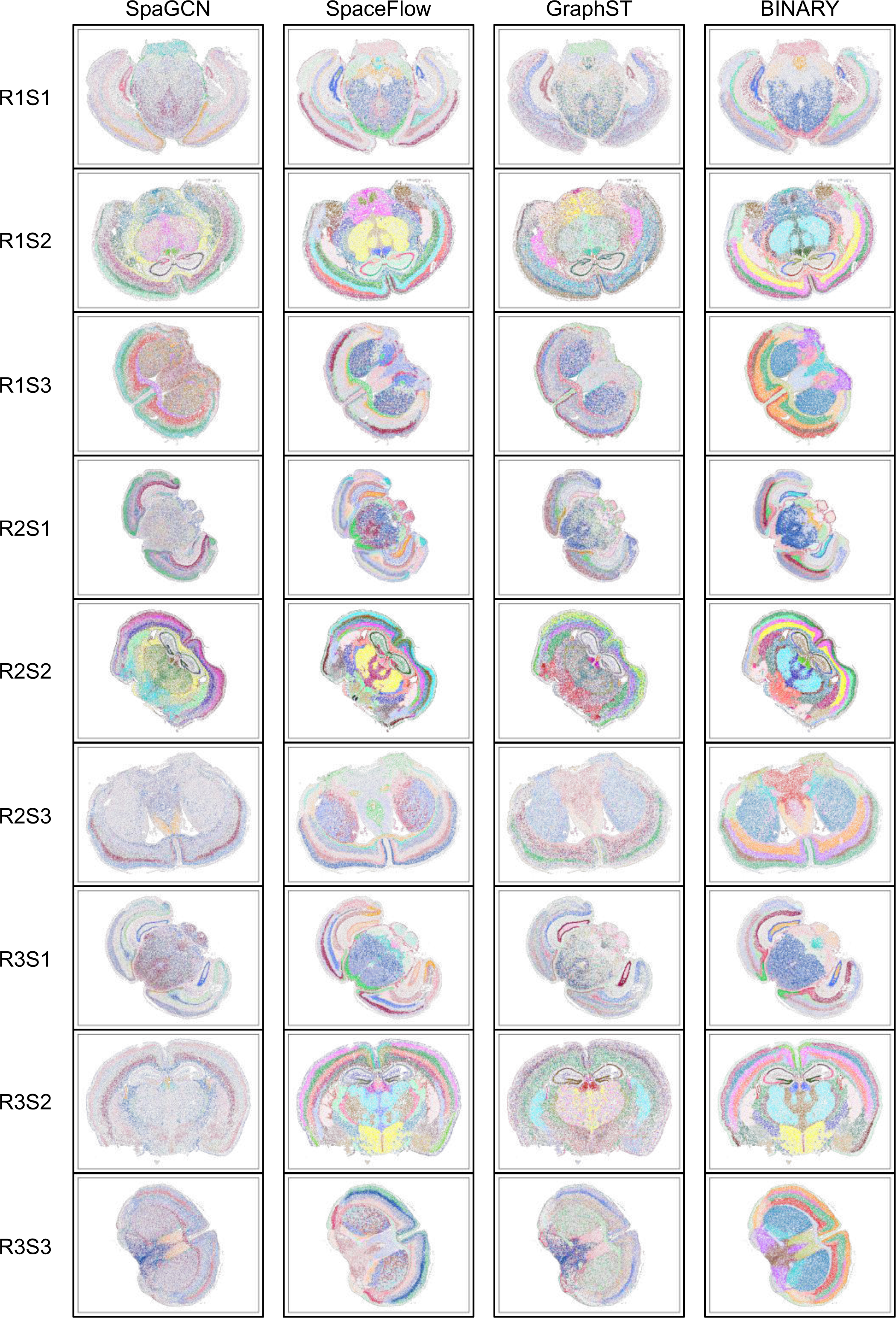
Comparative spatial domain identification of state-of-the-art spatial clustering methods on nine individual large-scale MERSCOPE datasets. Methods that support these large-scale MERSCOPE datasets include SpaGCN, SpaceFlow, GraphST, and BINARY. Notably, SEDR and STAGATE encounter memory overflow errors when running on one 80GB NVIDIA A100 GPU.

**Supplementary Fig. 8.**
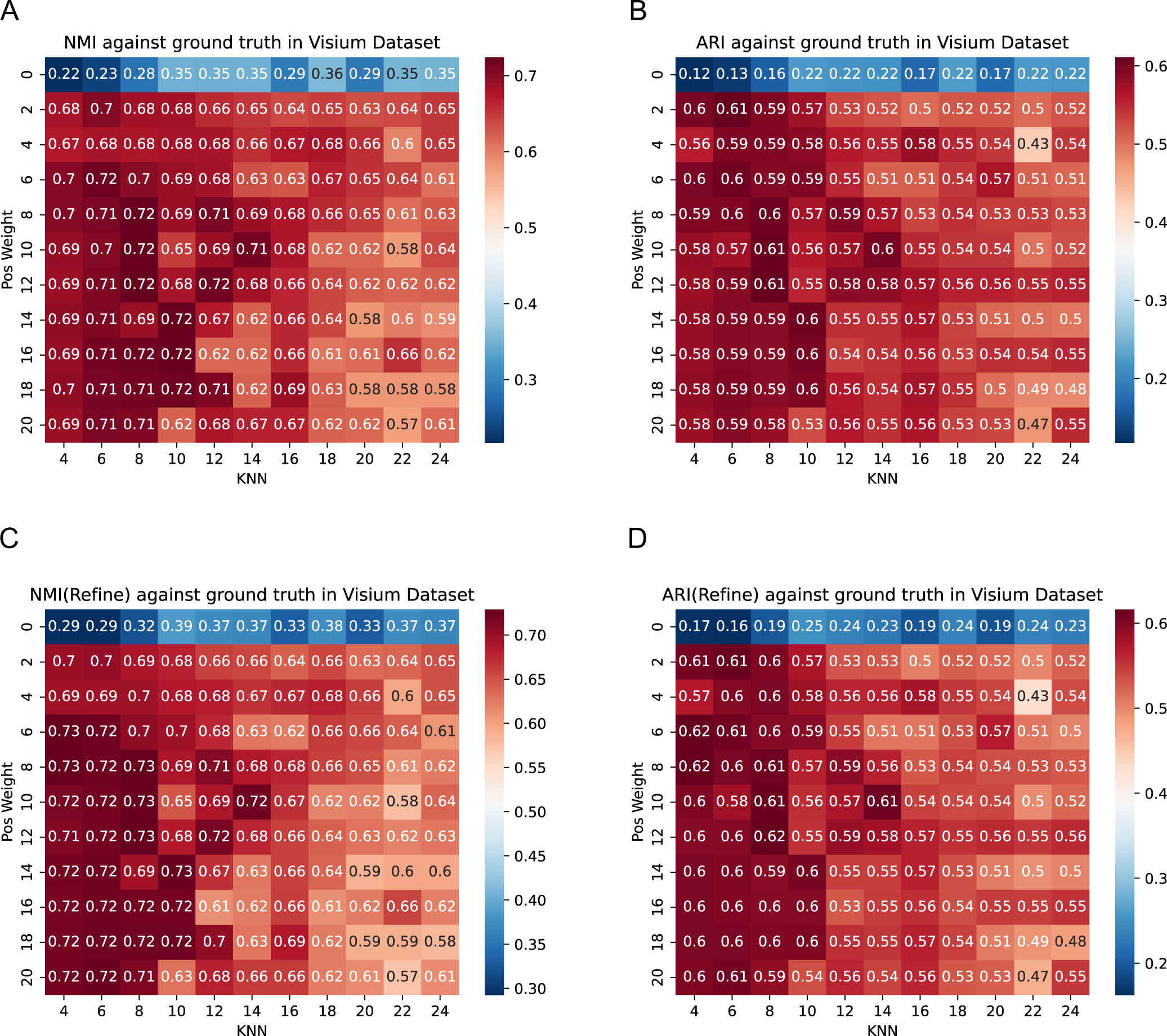
Sensitivity analysis of BINARY’s hyperparameters using the Visium dataset. **a,** Heatmap illustrating the Normalized Mutual Information (NMI) between BINARY clustering results and manual annotations (ground truths) on the Visium dataset across varying positive weight values (*w*_*pos*_) and KNN ranges (the ‘cutoff’ of knn). **b,** Heatmap showing the Adjusted Rand Index (ARI) between BINARY clustering results and manual annotations (ground truths) on the Visium dataset across different positive weight values (*w*_*pos*_) and KNN ranges (the ‘cutoff’ of knn). **c,** Heatmap depicting the NMI between refined BINARY clustering results post-processing and manual annotations (ground truths) on the Visium dataset for varying positive weight values (*w*_*pos*_) and KNN ranges (the ‘cutoff’ of knn). **d,** Heatmap presenting the ARI between refined BINARY clustering results post-processing and manual annotations (ground truths) on the Visium dataset for a range of positive weight values (*w*_*pos*_) and KNN settings (the ‘cutoff’ of knn). The x-axis represents the KNN values (the ‘cutoff’ of knn) used in graph construction, ranging from 4 to 24 in increments of 2. The y-axis represents the positive weight values (*w*_*pos*_), varying from 0 to 20 in steps of 2. Analyses are based on slice151673 of the DLPFC dataset. Each experiment was executed ten times, and the best result is reported.

**Supplementary Fig. 9.**
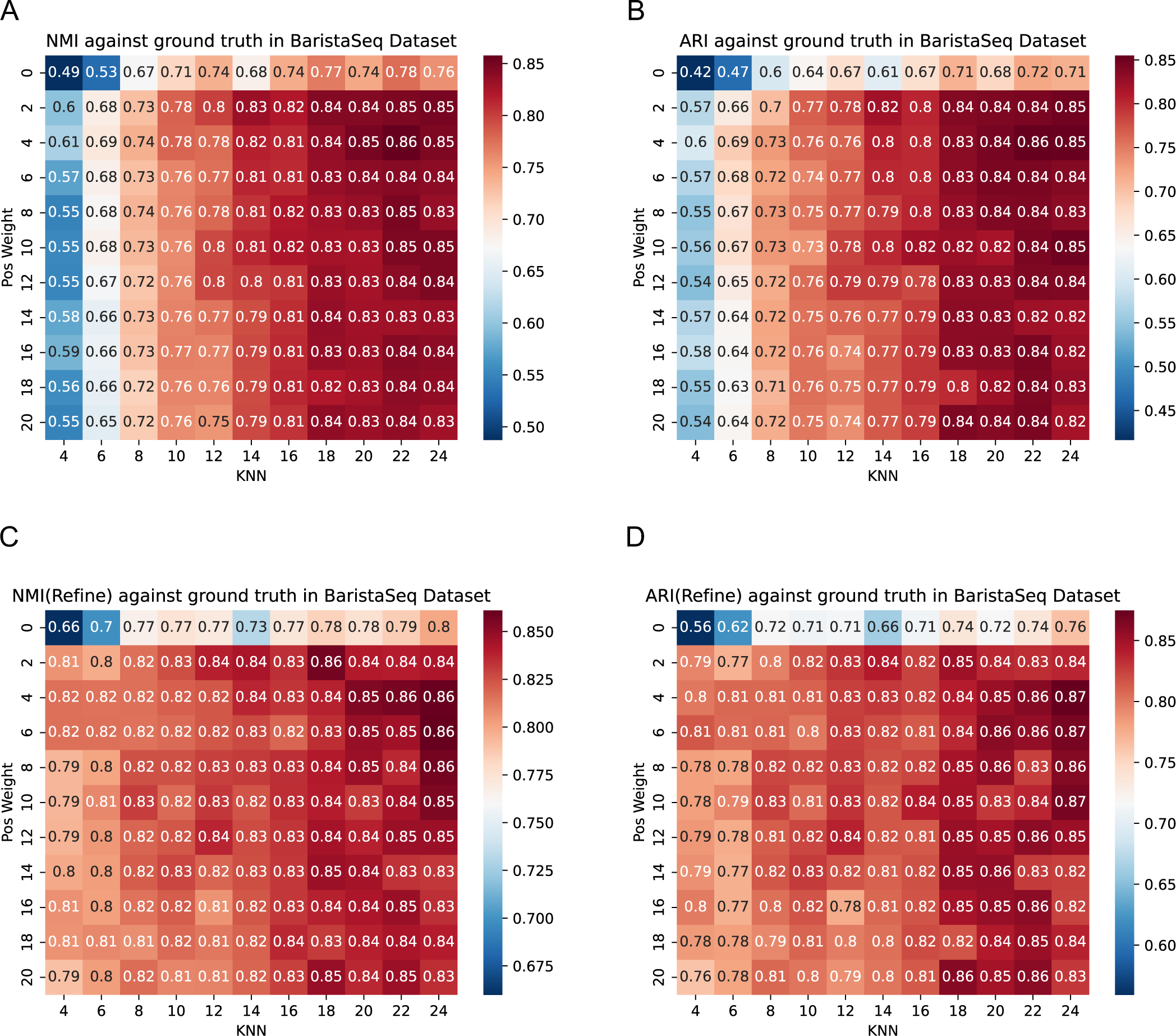
Sensitivity analysis of BINARY’s hyperparameters on the BaristaSeq dataset. **a,** Heatmap of the Normalized Mutual Information (NMI) between the clustering results of BINARY on the BaristaSeq Dataset and the ground truths (Manual annotations) at varying positive weight (*w*_*pos*_) and KNN range (the ‘cutoff’ of knn). **b,** Heatmap of the Adjusted Rand Index (ARI) between the clustering results of BINARY on the BaristaSeq Dataset and the ground truths (Manual annotations) at different positive weight (*w*_*pos*_) and KNN range (the ‘cutoff’ of knn). **c,** Heatmap of the NMI between the refined clustering results of BINARY on the BaristaSeq Dataset and the ground truths (Manual annotations) at various positive weight (*w*_*pos*_) and KNN range (the ‘cutoff’ of knn). **d,** Heatmap of the ARI between the refined clustering results of BINARY on the BaristaSeq Dataset and the ground truths (Manual annotations) across different positive weight (*w*_*pos*_) and KNN range (the ‘cutoff’ of knn). The x-axis represents the KNN values (the ‘cutoff’ of knn) utilized during graph construction, ranging from 4 to 24 in increments of 2. The y-axis denotes the positive value weights (*w*_*pos*_), ranging from 0 to 20 in increments of 2. The presented results are based on slice 2 of the BaristaSeq Dataset. Each experimental condition was replicated tenfold, and only the optimal outcomes were documented.

**Supplementary Fig. 10.**
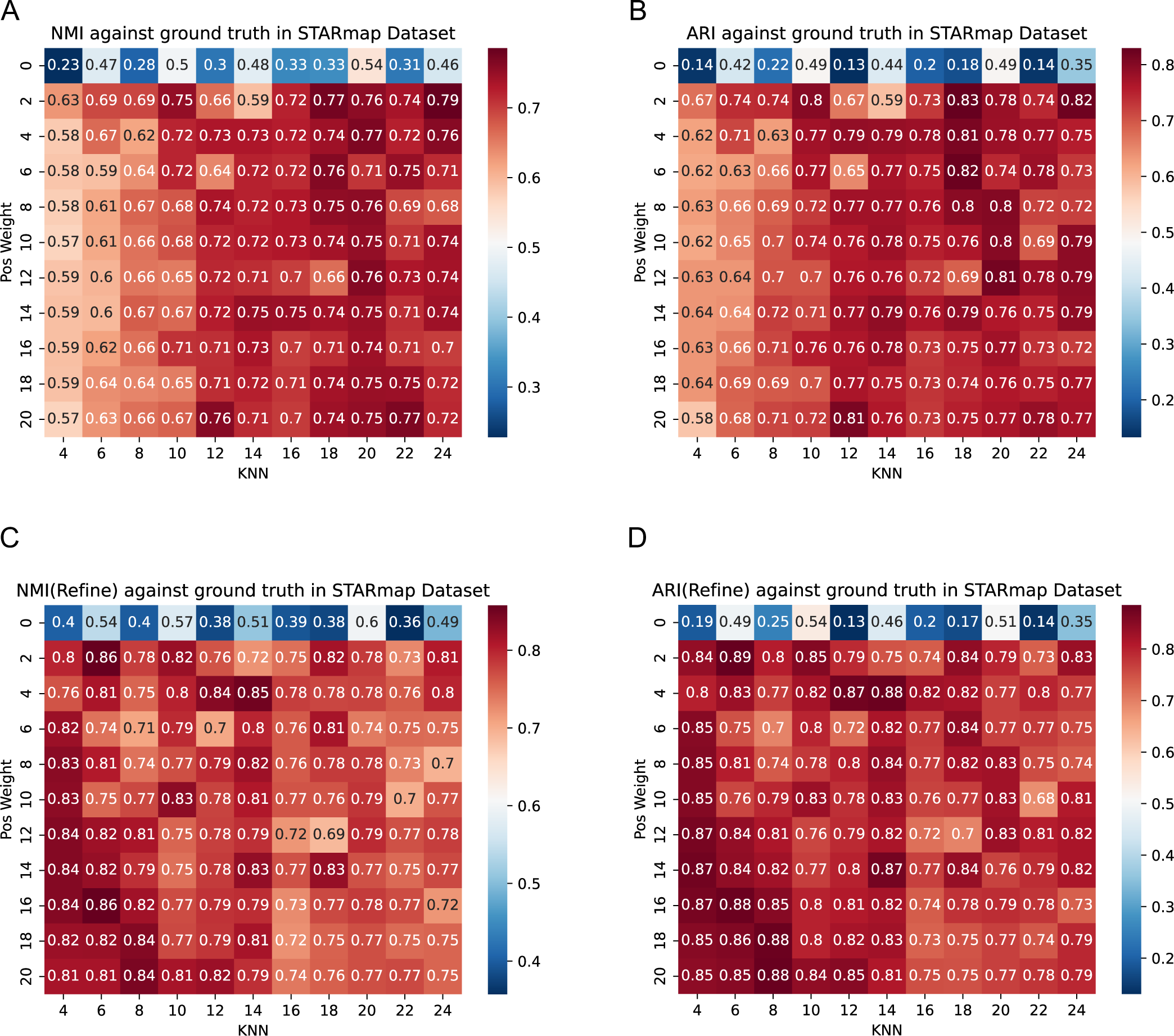
Sensitivity analysis of BINARY’s hyperparameters with the STARmap dataset. a, Heatmap of the Normalized Mutual Information (NMI) between BINARY’s clustering results and ground truths (Manual annotations) on the STARmap dataset., under varying positive value weights (*w*_*pos*_) and KNN (the ‘cutoff’ of knn). b, Heatmap of the Adjusted Rand Index (ARI) between BINARY’s clustering results and ground truths (Manual annotations) on the STARmap dataset., under varying positive value weights (*w*_*pos*_) and KNN (the ‘cutoff’ of knn). c, Heatmap of the NMI between BINARY’s refined clustering results and ground truths (Manual annotations) on the STARmap dataset., under varying positive value weights (*w*_*pos*_) and KNN (the ‘cutoff’ of knn). d, Heatmap of the ARI between BINARY’s refined clustering results and ground truths (Manual annotations) on the STARmap dataset., under varying positive value weights (*w*_*pos*_) and KNN (the ‘cutoff’ of knn). The x-axis represents the KNN values (the ‘cutoff’ of knn) used for graph construction, ranging from 4 to 24, with an increment of 2. The y-axis indicates the positive weights (*w*_*pos*_), spanning from 0 to 20, with an increment of 2. Analyses were based on BZ5 of the STARmap dataset. Each experimental condition was replicated ten times, and the optimal outcome from these replicates was selected for presentation.

**Supplementary Fig. 11.**
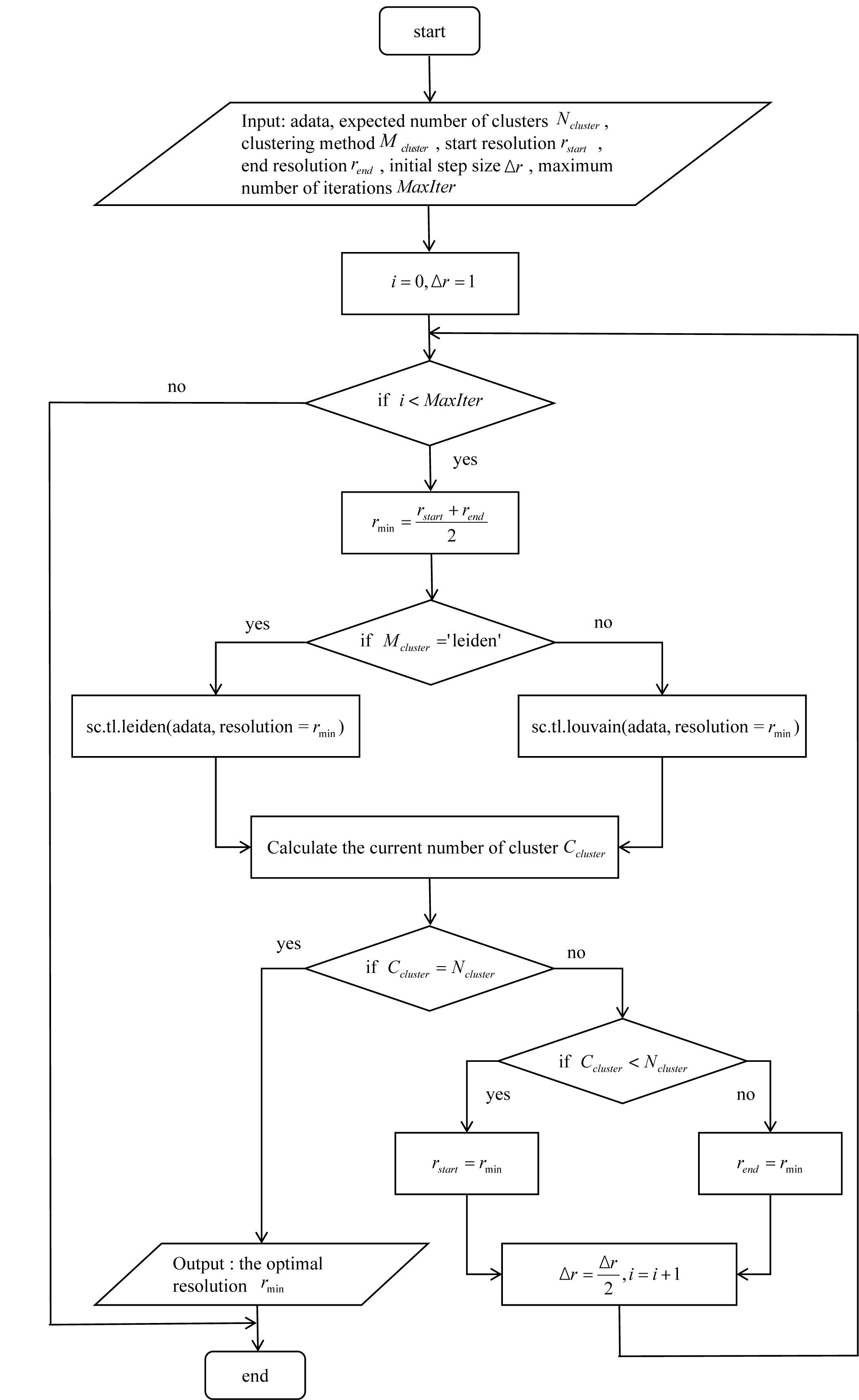
Binary search-based optimal resolution method for community detection clustering on BINARY. For the Louvain or Leiden spatial clustering methods implemented by the SCANPY package, BINARY introduces an iterative method to find the optimal resolution through binary search. The essence of this approach lies in the swift exploration of different resolution values using binary search, dynamically adjusting the step size until the desired number of clusters for the given data is achieved. This method allows for a rapid and precise determination of the resolution corresponding to a predefined cluster number.

**Supplementary Fig. 12.**
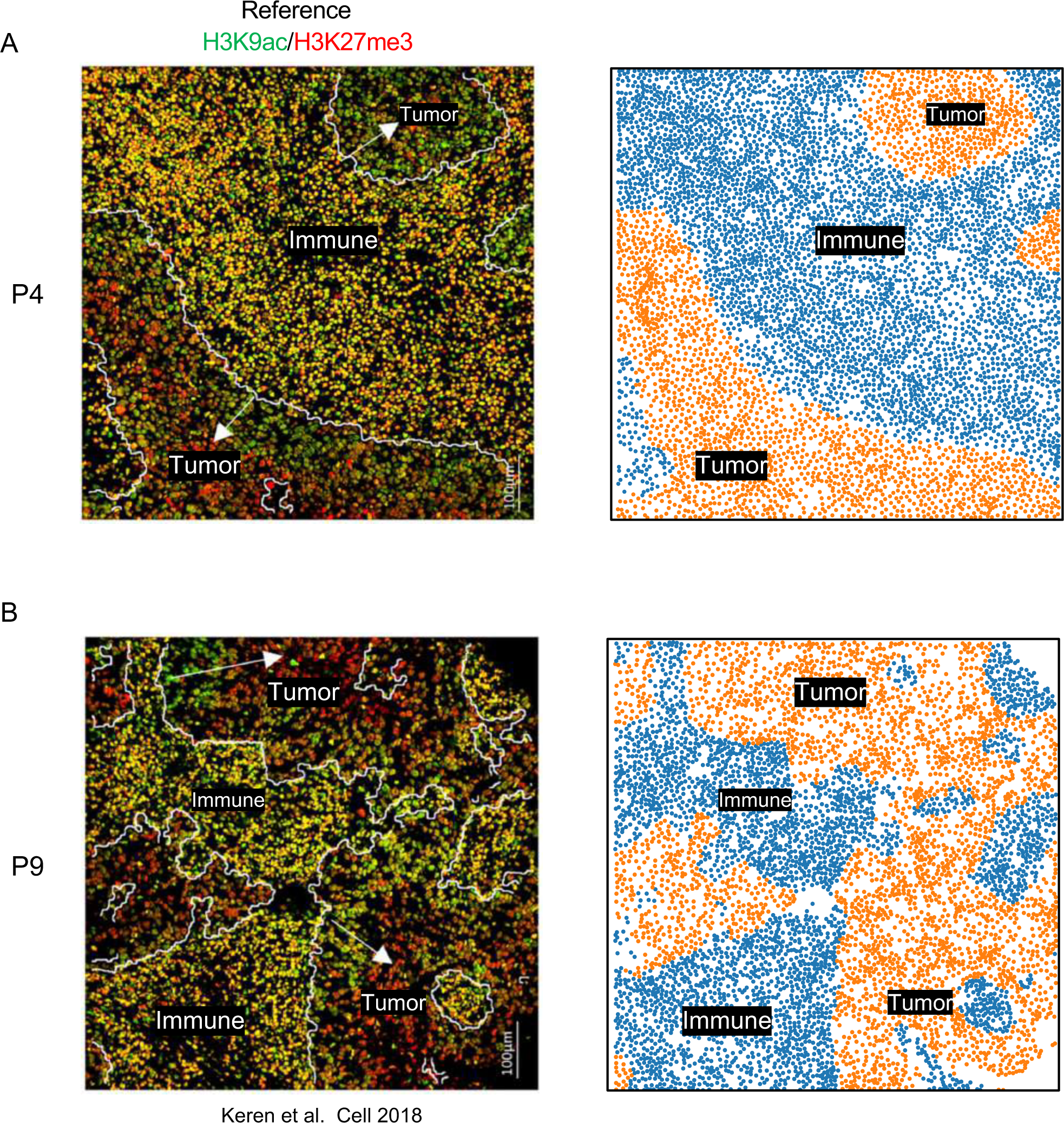
BINAYR demonstrates compatibility with spatial proteomics data. **a,** Clustering results from BINAYR on spatial proteomics data from patient 4, generated by MIBI-TOF (Right), align consistently with the reference (Left). **b,** Clustering outcomes by BINAYR on spatial proteomics data from patient 9, produced using MIBI-TOF (Right), closely match the reference (Left).

**Supplementary Fig. 13.**
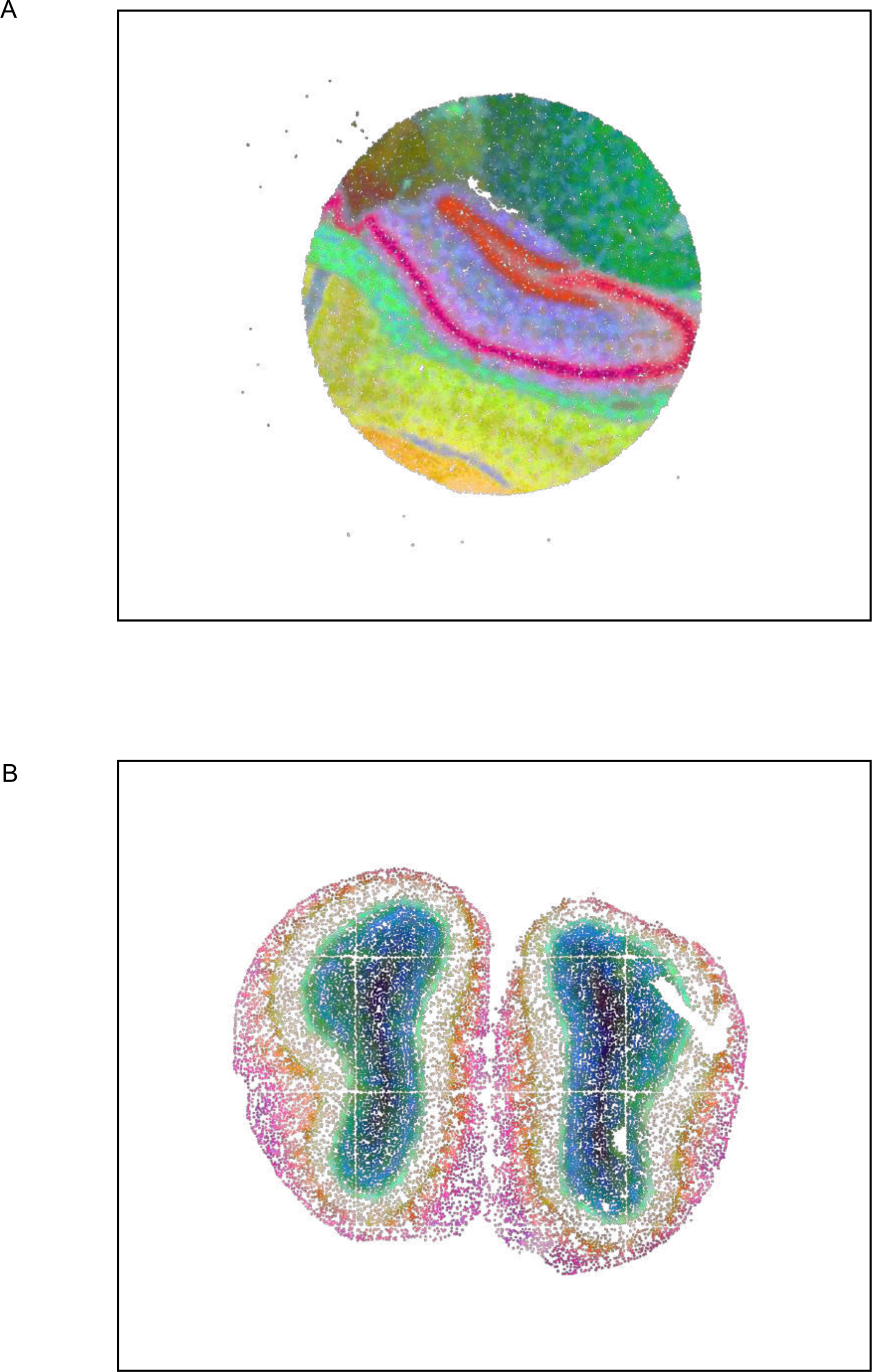
Depicting tissue continuity via embeddings derived from BINAYR. **a,** Tissue continuity representation in the Slide-seq dataset as portrayed by SOView. **b,** Illustration of tissue continuity within the Stereo-seq dataset through SOView.

**Supplementary Fig. 14.**
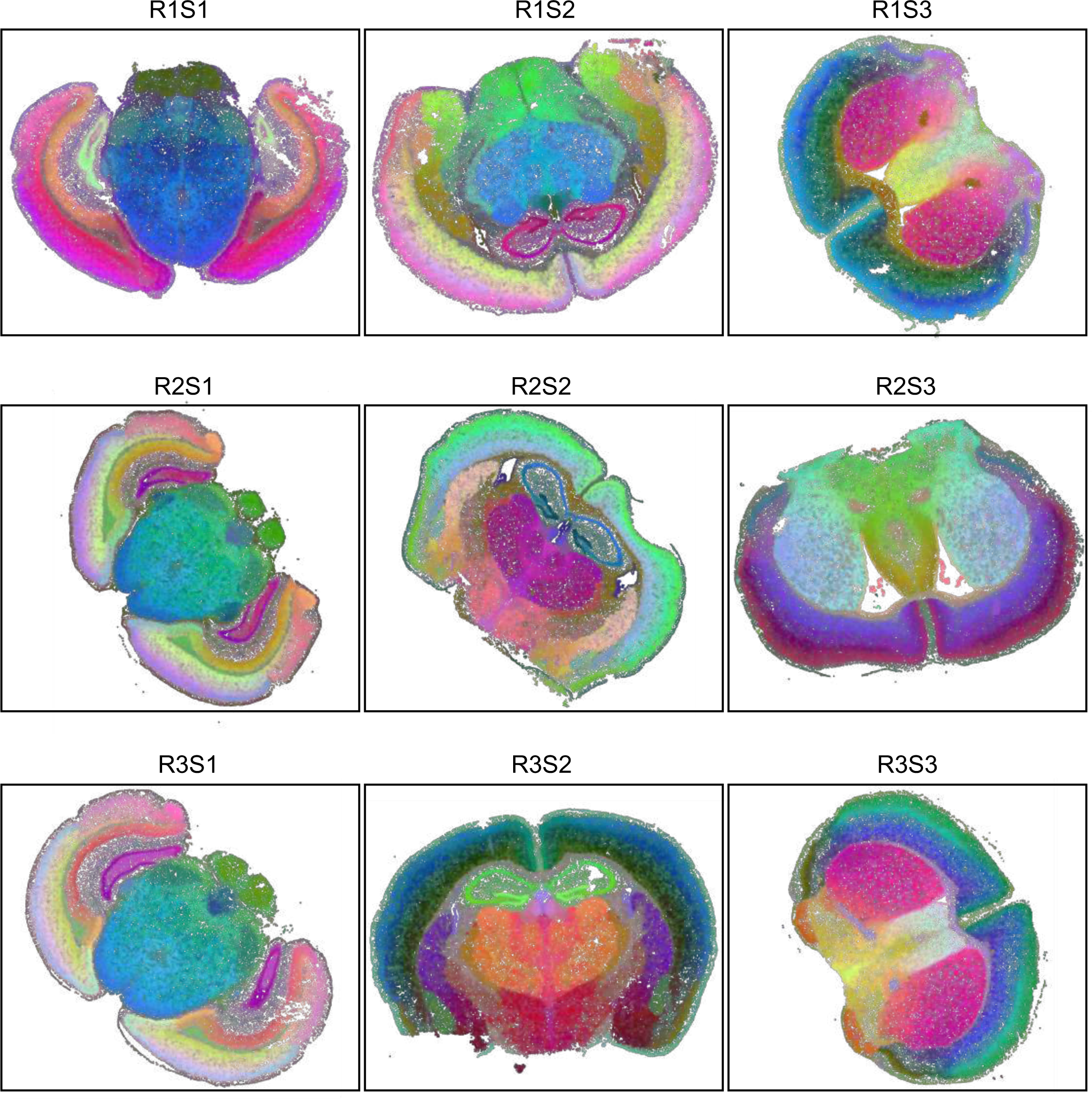
Visualization of tissue continuity by leveraging embeddings from BINAYR on a large-scale MERSCOPE dataset.

